# Enhancing sensitivity and versatility of Tn5-based single cell omics

**DOI:** 10.1101/2023.07.13.548833

**Authors:** Isabelle Seufert, Pooja Sant, Katharina Bauer, Afzal P. Syed, Karsten Rippe, Jan-Philipp Mallm

## Abstract

The analysis of chromatin features in single cells centers around Tn5 transposase and exploits its activity to simultaneously fragment target DNA and integrate adapter sequences of choice. This reaction provides a direct readout in the assay for transposase-accessible chromatin in single cells (scATAC-seq) to map open chromatin regions. However, a current limitation is the sparse coverage of open chromatin loci that are detected in a given single cell by droplet-based methods. Thus, enhancing Tn5 activity to improve genomic coverage of scATAC-seq or facilitating multi-omics readouts of chromatin features via Tn5 together with the transcriptome is of great interest. Here, we address these issues by optimizing scATAC-seq for an increased number of integrations per cell. In addition, we provide a protocol that combines mapping of histone modification with scRNA-seq from the same cell by targeting Tn5 to antibody-bound chromatin epitopes. Our experimental workflows improve the results obtained from the downstream data analysis and serve to better resolve epigenetic heterogeneity and transcription regulation in single cells.

## Introduction

For the analysis of chromatin features in single cells, Tn5 transposase is a crucial enzyme as it can simultaneously fragment target DNA across the genome and integrate adapter sequences of choice (Adey, 2021). This “cut-and-paste” reaction is referred to as tagmentation and has been initially used for preparation of DNA sequencing library. Usage of hyperactive Tn5 enzymes (Picelli et al., 2014) makes tagmentation an efficient process that can be conducted *in situ*, which is a prerequisite for its application in single cell sequencing (sc-seq). The direct application of tagmentation to analyze chromatin is the assay for transposase-accessible chromatin in single cells (scATAC-seq) (Buenrostro et al., 2015;Klemm et al., 2019). It maps enhancers and promoters that are in an open and *bona fide* active state and thus provides valuable information on transcription regulation mechanism and cellular heterogeneity with respect to cis-regulatory elements (Jiang et al., 2023). Furthermore, Tn5 is used in sc-seq multiomics readouts that yield, for example, scATAC-seq and scRNA-seq data from the same cell, albeit frequently with a lower sensitivity as compared to applying the two readouts in separate experiments (Dimitriu et al., 2022). Other types of single cell multiome analysis (e.g., RNA with CUT&Tag-seq of histone modifications) require specific modifications of the Tn5 mediated tagmentation reaction like using specific adapters loaded on a Protein A-tagged Tn5 in a so-called cut-and-tag reaction (Lahnemann et al., 2020;Lee et al., 2020). For all these applications, improvements to the Tn5 tagmentation reaction would advance the single cell analysis, which is frequently limited by data sparsity. For example, a typical coverage in scATAC-seq experiments with the Chromium drop-seq platform from 10x Genomics is 7,000 accessible sites per cell out of >100,000 sites that are detected by (pseudo-)bulk ATAC-seq (Li et al., 2021). Thus, the number of open loci detected in a given cell is rather sparse, which limits the resolution of cellular heterogeneity based on the ATAC signal. Aggregation of similar cells is one approach to address this issue (Pliner et al., 2018) but this leads to the loss of information on stochastic differences of chromatin accessibility between individual cells. Likewise, there is also a need to improve the sensitivity for the chromatin readout in multiomics experiments that map the transcriptome in the same cell together with ATAC or another chromatin feature via CUT&Tag (C&T) approaches.

Here, we optimized Tn5 activity to increase the sensitivity of scATAC-seq and described how its adapter loading can be made compatible with existing scATAC and multiome workflows for which the adapter configuration is unknown. We demonstrate that our protocols improve data quality and reduce sparsity without affecting scRNA-seq data quality for a more versatile multiome analysis.

## Results

We developed optimized experimental workflows that use differently loaded Tn5 preparations to improve Tn5-based analyses of single cells (**Fig. 1A**). Our work comprised three different reaction types: (i) In a protocol termed scTurboATAC we optimized the detection of open chromatin in scATAC-seq experiments over the standard protocol used with the Chromium platform from 10x Genomics. (ii) For the multiome protocol that combines scRNA-seq and scATAC-seq from the same cell (scMultiome-ATAC) we provide a protocol that uses phosphorylated adapters. (iii) The application for scC&T-seq together with scRNA-seq (scMultiome-C&T) involves Tn5 tagged with Protein A to target the enzyme in situ to the chromatin bound antibody. For this type of sc-seq analysis, we described a multiome analysis where histone H3 lysine 27 trimethylation is mapped alongside gene expression. Subsequent to the Tn5 reaction, the downstream processing followed standard protocols that resulted in efficient library preparation for the three applications described here (**Fig. 1B**).

**Fig. 1.**
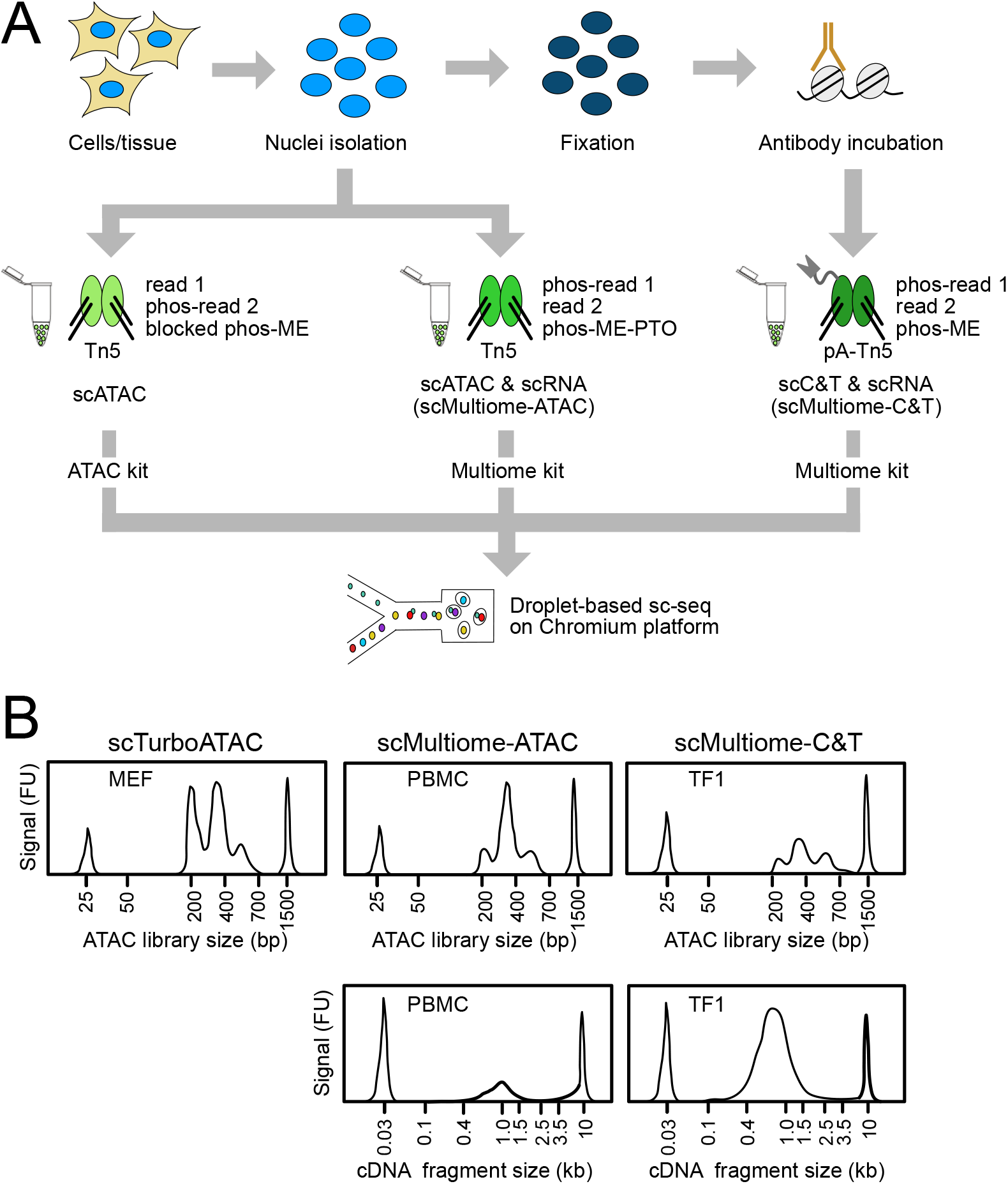
Experimental workflow. (**A**) Experimental procedure and use of the differently loaded Tn5 preparations. For scATAC and the multiome workflow nuclei were prepared and incubated with Tn5. Importantly, adapters varied between the protocols. For scMultiome-ATAC we used phosphorylated adapters, while the Tn5 for scMultiome-C&T was tagged with Protein A to facilitate binding to the antibody against a given chromatin feature. Following the Tn5 reaction, the downstream processing follows standard protocols. (**B**) Exemplary tapestation profiles are displayed for either the scATAC-seq or scC&T-seq libraries and corresponding cDNA fragments for the multiome protocols. MEFs and TF1 cells were directly used from cell culture while PBMCs were obtained from viably frozen aliquots. All profiles shown were generated with the Tn5-H100 enzyme preparation.

### scTurboATAC increases the number of open sites detected

By varying experimental parameters, we found that the activity of Tn5 is a limiting factor for scATAC-seq. We tested the activity of different concentrations of an in-house Tn5 preparation (Tn5-H) loaded with the adapters listed in **Table 1**. Its activity was compared in a qPCR assay to Tn5 enzymes from 10x Genomics (Tn5-TXG) as well as the Illumina TDE1 enzyme (Tn5-ILMN) by qPCR (**Fig. S1**).

**Table 1.**
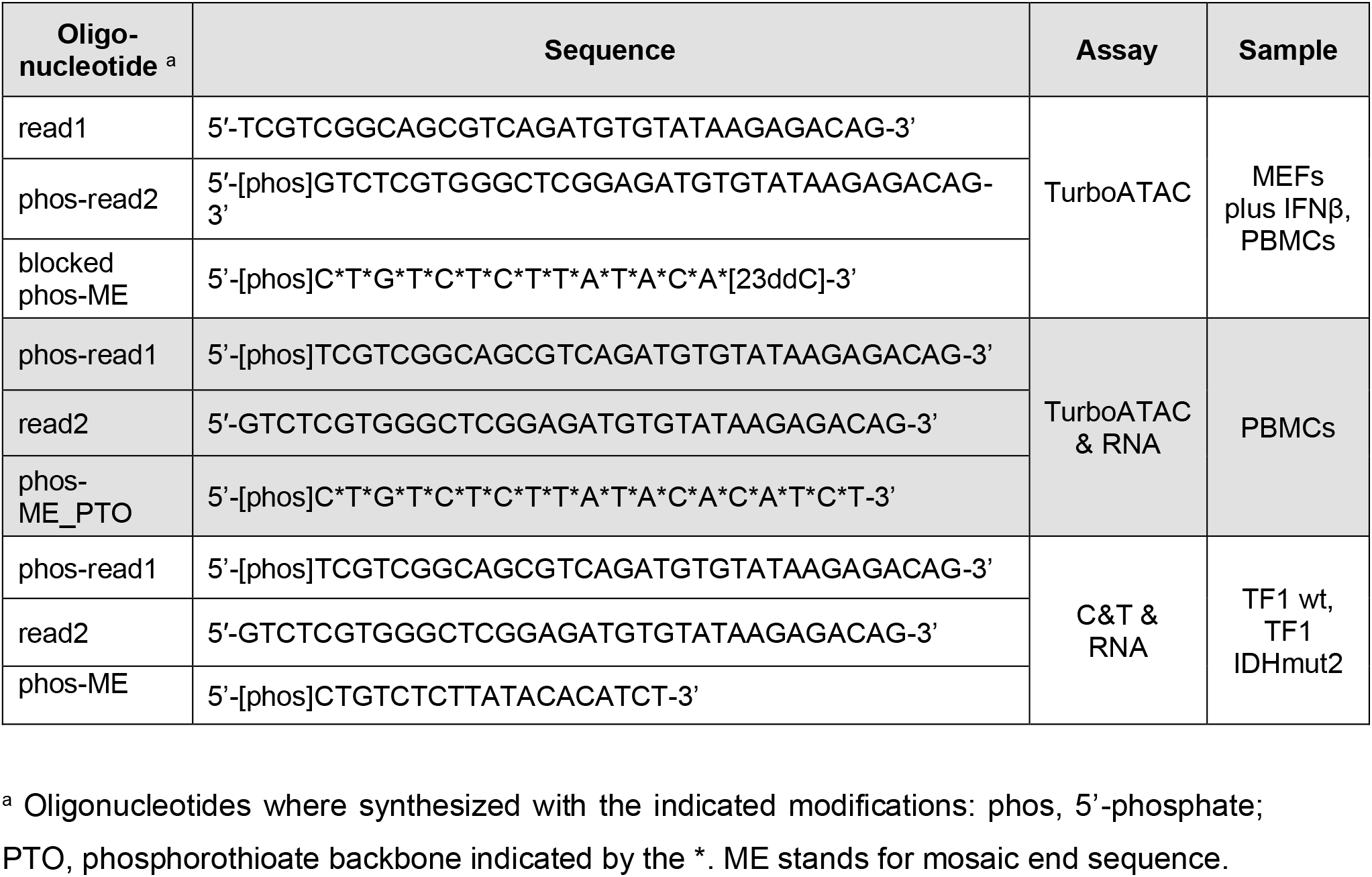
Oligonucleotides used for Tn5 loading.

The highest concentration of in-house Tn5 (Tn5-H100) that we used showed a four-fold higher activity per volume in terms of product yield as compared to the Tn5 enzyme provided in the 10x Genomics scATAC-seq version 2 (v2) kits (Tn5-TXGv2) and an about 1.3-fold higher activity per volume than Tn5-TXGv1.1 (**Fig. S1A**). We also tested different buffer compositions and found that the buffer provided in the 10x Genomics kit performed best. The Tn5-ILMN enzyme activity was comparable to Tn5-TXGv1.1 (**Fig. S1A, B**).

We then compared our Tn5-H100 and Tn5-H30 preparations to the Tn5-TXGv1.1 enzyme using mouse embryonic fibroblasts (MEFs) treated with interferon β (IFNβ) for 6 hours. We found that Tn5-H100 yielded the best results with respect to different QC parameters (**Fig. 2A, Fig. S2A**): Recovered cell barcodes showed (i) higher TSS enrichment scores, (ii) higher number of unique fragments, and (iii) smaller fragments. In the following, we refer to the use of Tn5-H100 with the buffers provided by 10x Genomics as scTurboATAC vs scATAC for the use with Tn5-TXG.

**Fig. 2.**
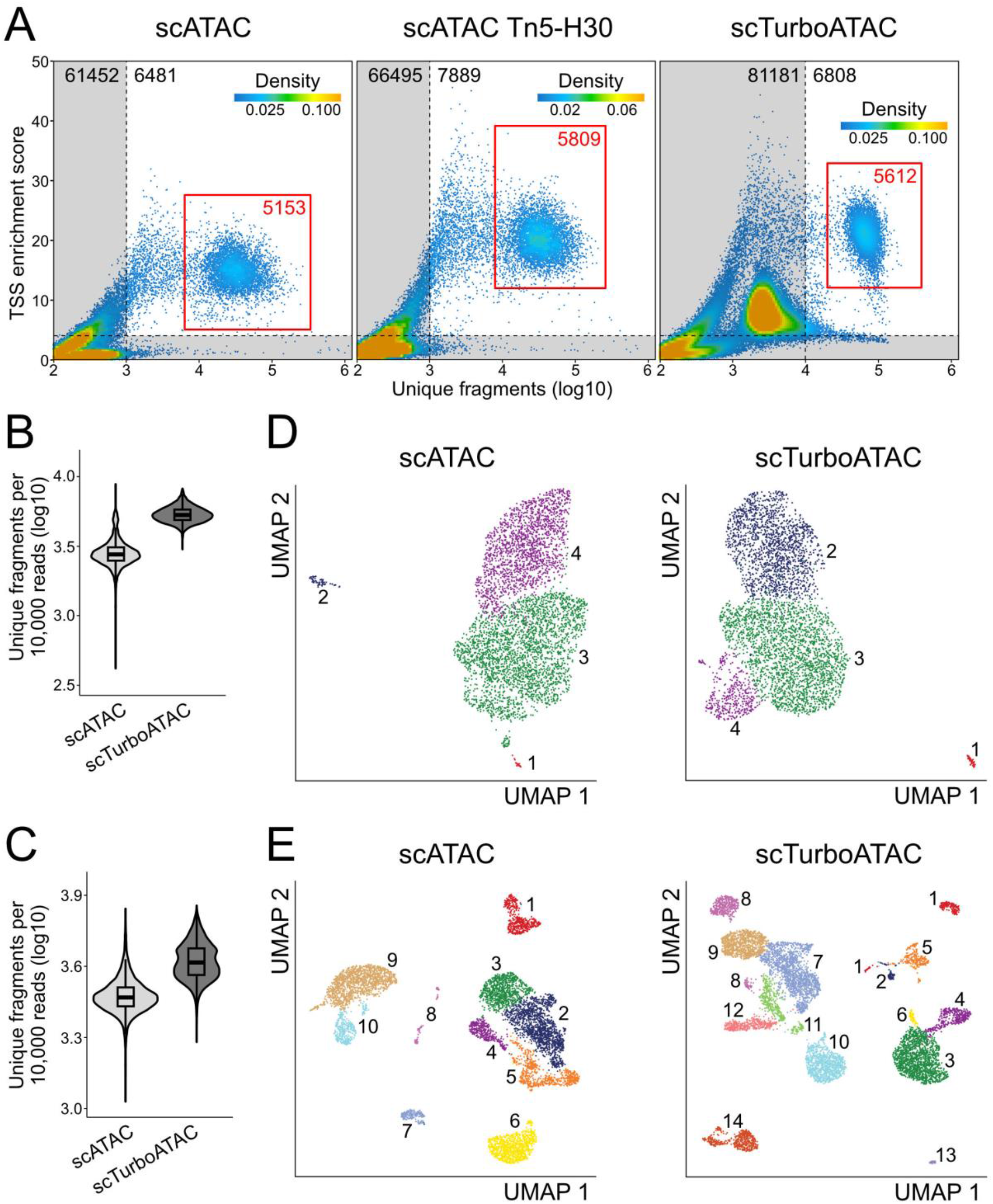
Quality assessment of scTurboATAC data. (**A**) TSS enrichment score over log10 number of unique fragments of cell barcodes in scATAC (left), scATAC with Tn5-H30 (middle) and scTurboATAC (right) of 6h IFNβ-stimulated MEFs. Colors of points reflect density of cell barcodes. Grey area marks low-quality cell barcodes. Red rectangle marks selected high-quality cells. (**B**) Number of unique fragments per 10,000 raw reads (log10) in scATAC and scTurboATAC of 6h IFNβ-stimulated MEFs. (**C**) Same as panel B for scATAC and scTurboATAC of PBMCs. (**D**) UMAP embedding of 6h IFNβ-stimulated MEFs from scATAC (left) and scTurboATAC (right). Each point represents one cell and is colored according to k-nearest neighbor cluster. (**E**) Same as panel D for PBMCs.

For further analysis, we selected high-quality cell populations of 5,000 to 6,000 cells for the three samples (**Fig. 2A**, red rectangle). The scTurboATAC protocol yielded an about 2-fold higher number of unique fragments normalized to sequencing depth than cells from the regular scATAC workflow (**Fig. 2B**). Similar to the mouse data, a higher number of unique fragments (absolute and normalized to sequencing depth) was detected in cryopreserved peripheral blood mononuclear cells (PBMC) as a test case when using Tn5-H100 as compared to Tn5-TXGv2 (**Fig. 2C, Fig. S2B**). Embedding and clustering of both data sets revealed and increased resolution of the scTurboATAC data for both MEF (**Fig. 2D**) and PBMC data (**Fig. 2E**). Using marker genes from our previous work (Muckenhuber et al., 2023) two distinct MEF subtypes (epithelial- and mesenchymal-like MEFs) were identified after removing minor clusters 1 and 2 (scATAC) and cluster 1 (scTurboATAC) that showed increased apoptosis module scores (**Fig. S2C**). Cluster 4 of scATAC and cluster 2 of scTurboATAC showed increased mesenchymal-like MEF marker module scores (**Fig. 2D, S3D**). The epithelial-like MEF markers were enriched in cluster 3 of scATAC and clusters 3 and 4 of scTurboATAC (**Fig. 2D, S3C**). Thus, the scTurboATAC protocol resolves two epithelial-like clusters, while only one cluster was detected with the standard scATAC protocol. Likewise, the PBMC data set displayed an increased number of clusters (14 instead of 10). These resolution improvements obtained with the scTurboATAC protocol are likely to reflect a higher complexity of the data that arises from the higher number of unique fragments.

### scTurboATAC improves the analysis of the transcription regulation

scTurboATAC was further evaluated for the analysis of transcription factor (TF) binding and gene regulatory mechanisms in MEFs stimulated with IFNβ for 6 hours. Calling peaks from the pseudo-bulk data yielded a higher number of accessible peaks per cell with the scTurboATAC protocol (**Fig. S3A, Fig. S3B**).

Furthermore, accessibility footprints at transcription start sites (TSSs) and at STAT1 and CTCF binding motifs showed higher enrichment in pseudo-bulk scTurboATAC than scATAC (**Fig. 3A**).

**Fig. 3.**
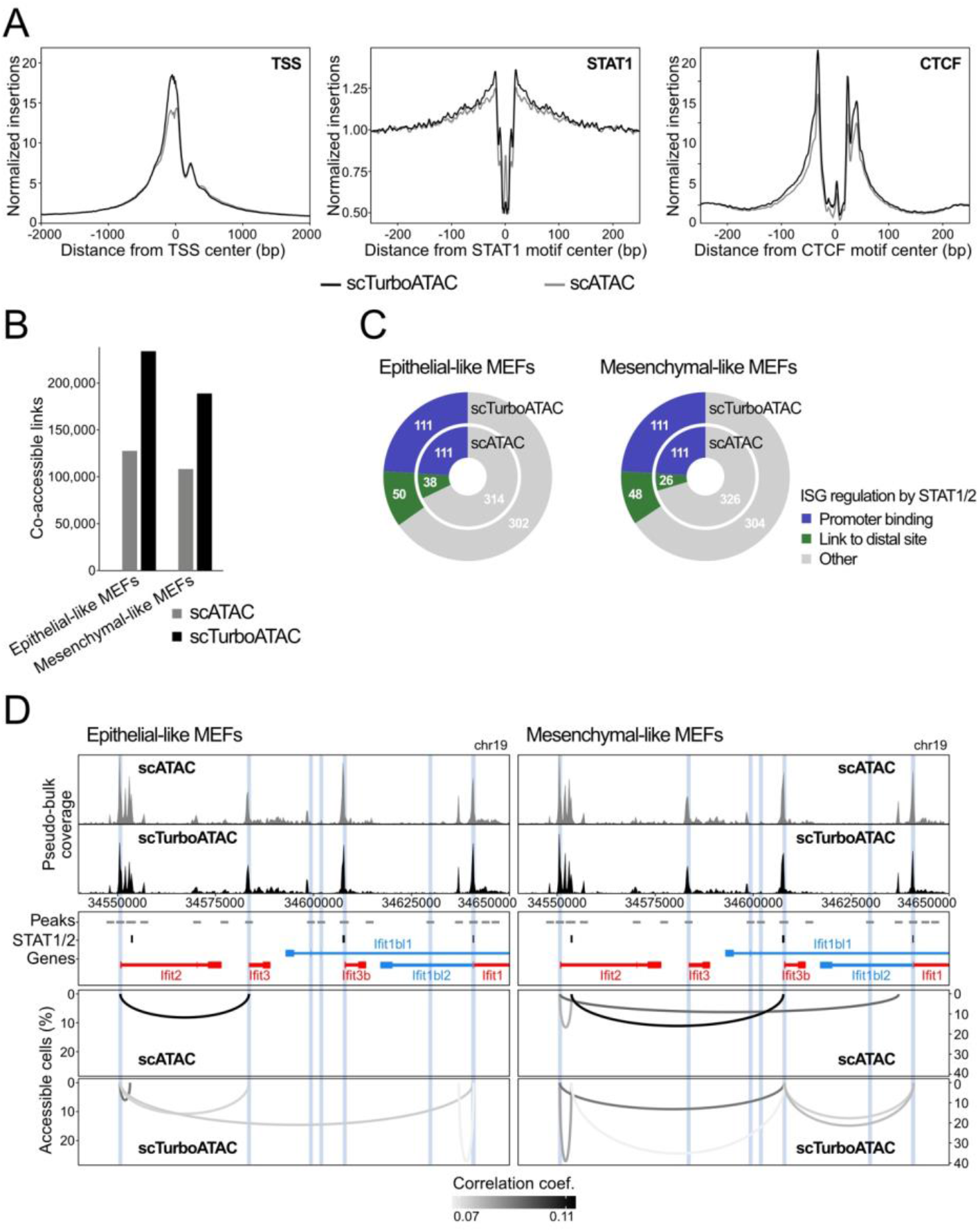
Comparison of scATAC and scTurboATAC. MEFs stimulated with IFNβ for 6 hours were analyzed. (**A**) Accessibility footprints at TSSs (left), STAT1 motifs (middle) and CTCF motifs (right) from scATAC and scTurboATAC. (**B**) Number of co-accessible links in scATAC and scTurboATAC of epithelial- and mesenchymal-like MEFs. (**C**) ISG regulation mechanisms according to STAT1/2 binding after IFNβ treatment in epithelial- (left) and mesenchymal-like MEFs (right). Promoter bound STAT1/2 (independent of the presence of additional links to distal STAT1/2 sites), blue; co-accessible link to a distal STAT1/2 peak, green; other ISGs, grey. (**D**) Co-accessibility in epithelial- (left) and mesenchymal-like MEFs (right) at the Ifit ISG cluster. Top: browser tracks of aggregated pseudo-bulk chromatin accessibility from single cells of scATAC and scTurboATAC. Middle: 2 kb regions around peaks from pseudo-bulk chromatin accessibility (grey); STAT1/2 peaks from bulk ChIP-seq (black); gene annotation (red, blue). 1 kb regions around ISG TSSs from bulk RNA-seq are marked in blue. Bottom: co-accessible links from scATAC and scTurboATAC.

Next, we applied our approach for co-accessibility analysis to MEF subtypes in scATAC and scTurboATAC using the RWire software as described previously (Muckenhuber et al., 2023). RWire reveals enhancer-promoter links from correlation between open sites of scATAC-seq data. For epithelial-like MEFs of scTurboATAC, only cluster 3 was considered. We observed twice as many co-accessible links above background co-accessibility and minimal percent accessible cells in epithelial-as well as mesenchymal-like MEFs of scTurboATAC than scATAC (**Fig. 3B**). Additionally, co-accessible links from scTurboATAC showed higher correlation coefficients and percent accessible cells than co-accessible links from scATAC (**Fig. S3E**). In our previous study, we observed that the transcription activation of ISGs is partly regulated by simultaneous STAT1 and STAT2 (STAT1/2) binding to ISG promoters as well as distal regulatory sites (Muckenhuber et al., 2023). We were able to link roughly 10 % of ISGs without STAT1/2 promoter binding to a distal STAT1/2 binding event by co-accessibility analysis of 6h IFNβ-stimulated epithelial-as well as mesenchymal-like MEFs from scTurboATAC (**Fig. 3C**). In contrast, only 5-8 % of ISGs were linked to a distal STAT1/2 binding event by co-accessibility analysis of 6h IFNβ-stimulated epithelial-as well as mesenchymal-like MEFs from scATAC. An example for the improved detection of ISG regulation by distal STAT1/2 binding events with scTurboATAC is shown for the *Ifit* gene cluster in **Fig. 3D**. The 150 kb locus depicted contains six ISGs (*Ifit1, Ifit1bl1, Ifit1bl2, Ifit2, Ifit3*, and *Ifit3b*) that were previously identified in MEFs (Muckenhuber et al., 2023). For *Ifit1, Ifit1bl2* and *Ifit3b* promoters, direct STAT1/2 binding was observed in IFNβ-stimulated MEFs. We detected three co-accessible links between ISG promoters and distal STAT1/2 binding events in epithelial- and mesenchymal-like MEFs of scATAC. All co-accessible links were also detected in epithelial- and mesenchymal-like MEFs of scTurboATAC, where they showed higher confidence by higher percent accessible cells. In addition, six co-accessible links between ISG promoters and STAT1/2 binding events were only detected with scTurboATAC. In mesenchymal-like MEFs of scATAC, we detected one unspecific link between the *Ifit2* promoter and a distal peak without a STAT1/2 binding event or other ISG promoter. This link was not observed in mesenchymal-like MEFs of scTurboATAC.

### scTurboATAC enhances the cell type annotation

Next, we compared scTurboATAC and scATAC with the Tn5-TXGv2 enzyme with respect to cell type resolution by clustering and marker gene activity (**Fig. 4A, Tab. S2**). The scTurboATAC protocol specifically improved the detection of progenitor cells, classical monocytes/dendritic cells and provided a higher resolution of the B cell cluster (**Fig. 4A, B**). We identified a distinct B cell cluster in scTurboATAC (cluster 1) that was not resolved in scATAC data by integrating B cells from scATAC and scTurboATAC (**Fig. S4B**). When investigating the B cell clusters further, we found differential TF activities for PU.1-IRF, OCT2 and ATF3 among others (**Fig. 4B, C, Fig. S4C**). While OCT2 has been observed in late state B cell differentiation (Corcoran et al., 1993), PU.1-IRF plays a role in earlier lymphoid maturation (Scott et al., 1994). This could account for their contrary activity scores. Elevated ATF3 activity was mainly found in cluster 1 of scTurboATAC data, potentially pointing to stress-induced activation of B cells to repress pro-apoptotic genes and cytokine production (Ku and Cheng, 2020).

**Fig. 4.**
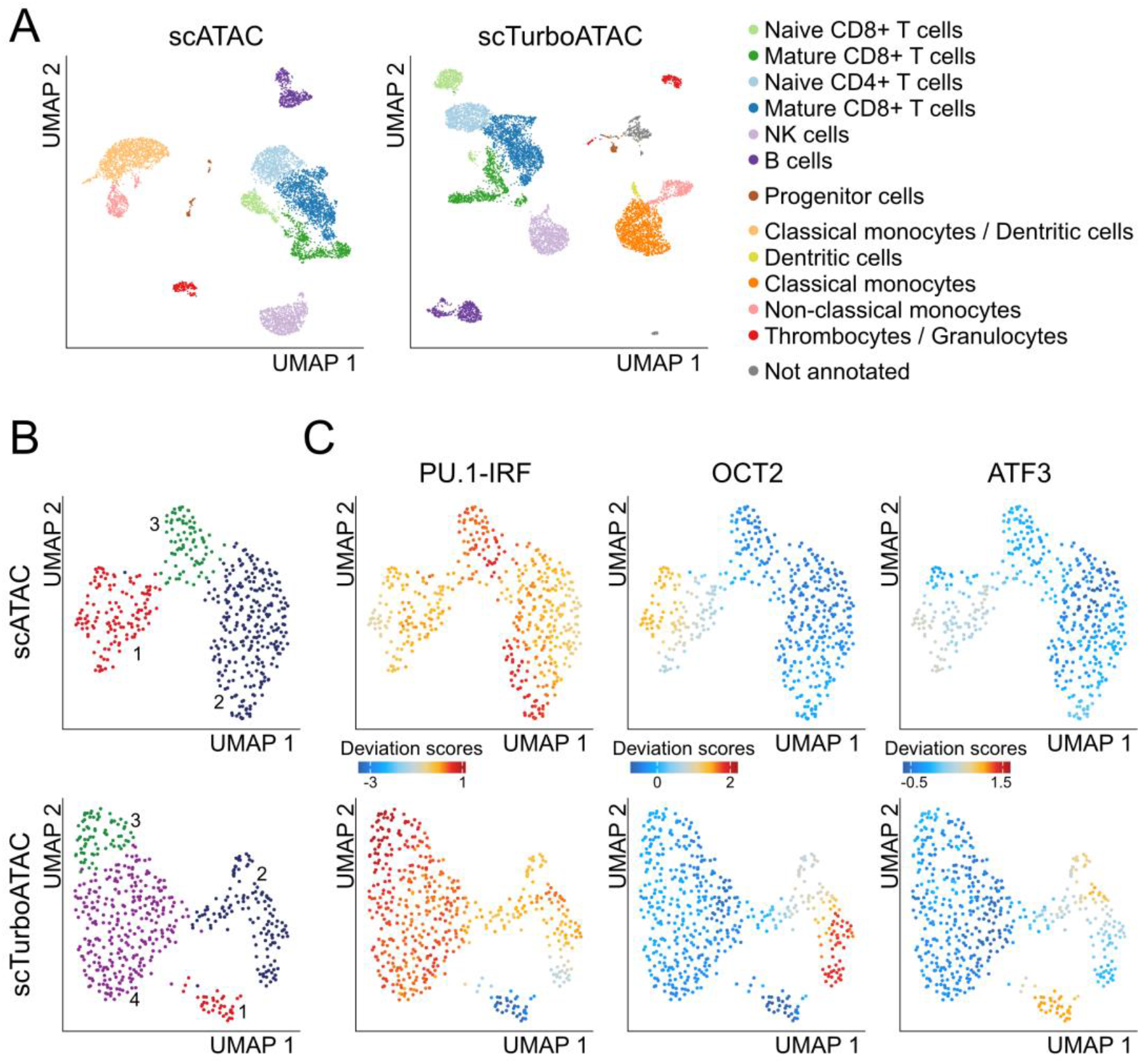
scTurboATAC resolves additional cell types in PBMCs. (**A**) UMAP embedding of PBMCs from scATAC (left) and scTurboATAC (right). Cells are colored according to annotated cell type by marker gene activity scores. (**B**) UMAP embedding of PBMC B cells from scATAC (top) and scTurboATAC (bottom). Cells are colored according to k-nearest neighbor clusters. (**C**) Same as panel B with cell coloring according to imputed transcription factor motif deviation scores.

### scTurboATAC can be integrated into an improved multi-omics analysis

Multi-omics approaches allow to directly link epigenetic heterogeneity with transcriptional profiles. We thus wanted to further exploit scTurboATAC for this type of application and adapted the loading of the transposase to be compatible with the 10x Genomics multiome chemistry. The latter utilizes a spacer oligonucleotide to add cell barcodes to the Tn5 fragments presumably by ligation. To account for this difference, we changed the oligonucleotides for loading (**Table 1**) and additionally used a Protein A-tagged Tn5 for a single-cell C&T-seq assay with RNA information from the same cell (**Fig. 1B**). To avoid self-dimerization for the multiome-ATAC workflow a complementary oligonucleotide with a phosphorothioate (PTO) backbone was used. This modification can be omitted for multiome-C&T due to the Tn5 capture and washing of unbound Tn5. Phosphorylation of Tn5_read1 was found to be essential to facilitate the barcode ligation step in the droplets.

In agreement with the scTurboATAC data, we observed a higher number of unique fragments normalized to sequencing depth when using our adapted Tn5-H100 in comparison to Tn5-TXGv2 with the multiome chemistry in PBMCs (**Fig. 5A left**) while the number of reads in TSSs was similar (**Fig. S5A, B**). UMI counts of transcripts normalized to sequencing depth showed the same yield in comparison to the standard workflow (**Fig. 5A right**). The number of clusters was higher for scMultiome-ATAC with Tn5-H100 (**Fig. 5B**) compared to the standard scMultiome-ATAC with Tn5-TXGv2 (**Fig. 5C**) for both for the ATAC readout alone as well as co-embedded ATAC and RNA data. When using only the gene expression data, the number of clusters remained the same (**Fig. 5B, C**). Thus, the improved ATAC reaction did not compromise the RNA readout in the scMultiome-ATAC protocol.

**Fig. 5.**
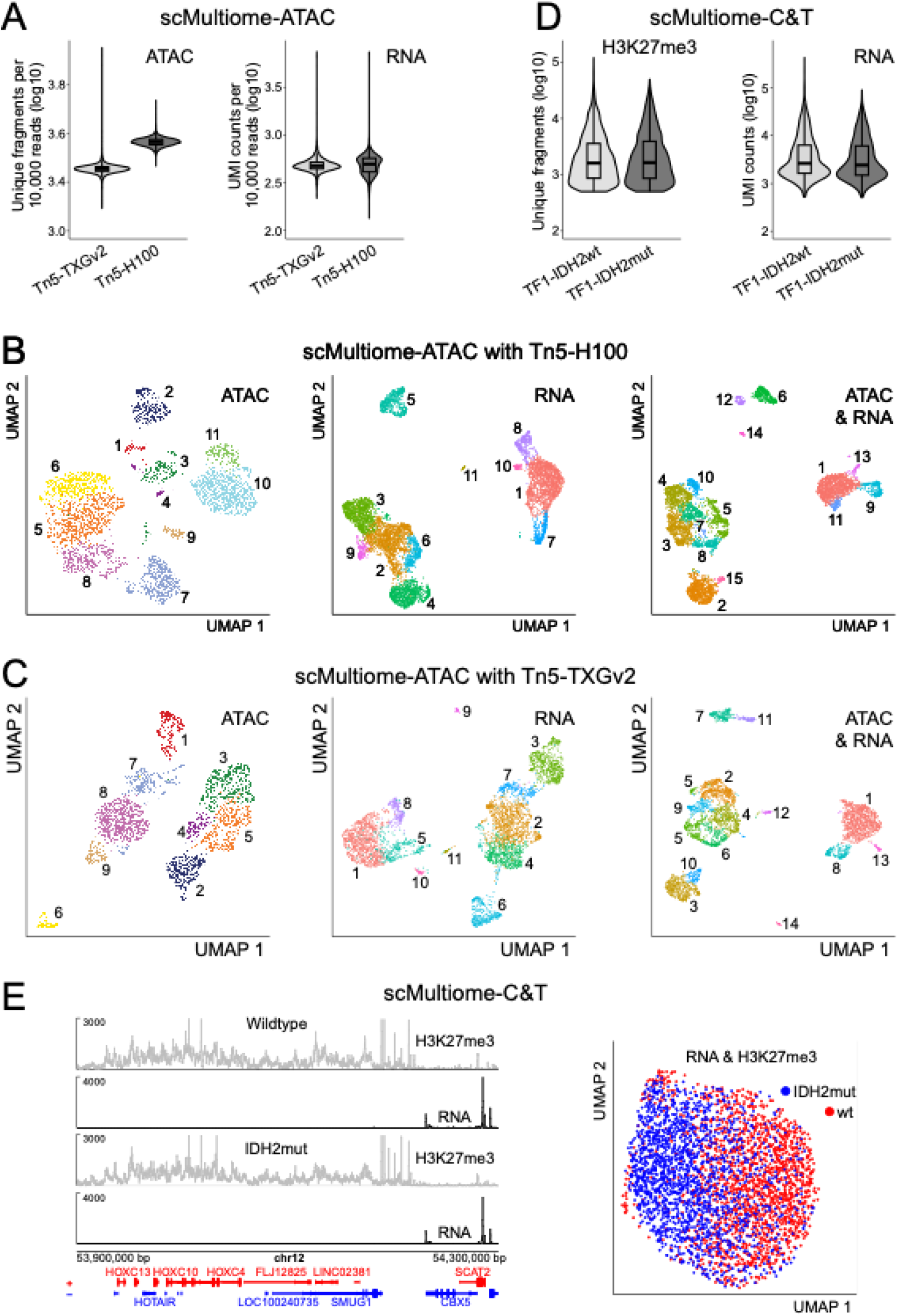
Multiome analysis. (**A**) scMultiome-ATAC analysis of PBMCs. The log10 number of unique fragments per 10,000 raw reads (ATAC, left) and UMI counts per 10,000 raw reads (RNA, right) is shown. (**B**) UMAP embedding of PBMCs from scMultiome-ATAC with Tn5-H100 based on ATAC (left), RNA (middle) and combined ATAC and RNA (right) readouts. Cells are colored according to k-nearest neighbor clusters. (**C**) Same as panel B but for the scMultiome-ATAC standard protocol with Tn5-TXGv2. (**D**) Analysis of TF1 wildtype and IDH2 mutated cell lines by scMultiome-C&T. The log10 number of unique fragments for H3K27me3 (left) and of UMI counts for RNA (right) are plotted. (**E**) scMultiome-C&T of TF1 wildtype and IDH2 mutated cells. Left, browser tracks of aggregated pseudo-bulk H3K27me3 (grey) and RNA (black) for an exemplary region with the genes on the plus (red) and minus (blue) strand shown at the bottom. Right, UMAP co-embedding of H3K27me3 and RNA. Cells are colored according to sample.

The scC&T analysis was performed in isogenic TF1 wildtype and mutant cell lines that carry the IDH2 R140Q mutation (Stein, 2016) with histone H3 trimethylation at lysine 27 (H3K27me3) as C&T target. Data from both samples show a similar yield in the number of H3K27me3 fragments (**Fig. 5D left**) with a bias for high-nucleosomal fragments (**Fig. S5C**), a high fraction of reads in peaks (**Fig. S5D**) and comparable UMI transcript counts (**Fig. 5D right**). Very broad regions of H3K27me3 enrichment were observed as depicted for an exemplary region of the HOXC gene cluster in **Fig. 5E left**. In both cell lines all genes within this H3K27me3-rich region were silenced while adjacent genes lacking H3K27me3 signal were highly expressed. UMAP of C&T and RNA data from both cell lines showed separate embedding with some overlap according to IDH2 mutation status (**Fig. 5E right**). Taken together, these data illustrate the versatility and specificity of the scTurboATAC and scMultiome-C&T techniques, which open up new ways for data analysis and experimental readouts.

## Discussion

Single-cell sequencing applications are frequently limited by data sparsity, which has been addressed both experimentally and bioinformatically (Lahnemann et al., 2020;Bouland et al., 2023;Sant et al., 2023). One strategy to counteract the sparsity for scATAC-seq data is to aggregate multiple similar cells (Pliner et al., 2018;Chen et al., 2019). However, this leads to the loss of information on stochastic differences between individual cells. The scTurboATAC protocol introduced here improved the experimental data acquisition and resulted in a higher library complexity with a consistently lower number of duplicates and higher number of integration events per cell (**Table S1**). These enhancements led to better results in the downstream analysis as shown for cell clustering and computation of TF motif footprints and activities. Furthermore, scTurboATAC improved the co-accessibility analysis to detect interactions between cis-regulatory elements, which was conducted here without aggregating cells to capture stochastic changes of chromatin accessibility.

Plate-based scATAC methods, similar to RNA analysis, could perform better than the droplet-based method used here (Xu et al., 2021). The droplet-based approaches, however, allow processing of several samples with thousands of cells each, which is needed to resolve cell types in complex tissues. Their throughput can be further improved by multiplexing samples combined with overloading the chip (Han et al., 2022). Thus, it will be interesting to compare the sensitivity of our scTurboATAC protocol to plated-based scATAC-seq methods and to evaluate its integration into multiplexing approaches.

As stated above, the relationship between enhancer activity, transcription factor binding and gene expression is of general interest (Fulco et al., 2019;Riegel et al., 2023;Uyehara and Apostolou, 2023). Techniques exist that allow the integration of separately acquired data types from the same sample. However, it is difficult to separate biological differences from confounding technical biases that need to be removed via the integration. Multiome readouts overcome this problem but frequently provide lower sensitivity (Dimitriu et al., 2022). Thus, improving sensitivity for each individual readout is crucial. To this end, we here also adapted and optimized loading of Tn5 to make it compatible with existing multiome workflows for which the adapter configuration is unknown. Furthermore, we demonstrate that our scMultiome-ATAC protocol improved data quality and reduced sparsity without affecting scRNA-seq data quality.

In comparison to the scMultiome-ATAC analysis, the combination of scRNA with scC&T-seq is more difficult to implement (Lahnemann et al., 2020;Lee et al., 2020), especially for repressive or promoter distal histone marks like H3K9me3 and H3K27me3. To address this point, we here also devised a scMultiome-C&T protocol with specific adapters loaded on a Protein A-tagged Tn5 that can be easily applied by other laboratories. A similar workflow has been described before but as a single readout without transcriptome information (Kaya-Okur et al., 2019). Here, we used our scMultiome-C&T method to analyze the repressive H3K27me3 histone modification together with the RNA from the same cell. An enrichment of H3K27me3 and repression of transcription was detected at loci that were previously described as silenced by H3K27me3 (Soshnikova and Duboule, 2009;Noordermeer et al., 2011). We expect that our technique will work well for all C&T-grade antibodies and can thus easily be adapted to other histone modifications of interest.

In summary, the scTurboATAC, scMultiome-ATAC and scMultiome-C&T protocols described here enhance the sensitivity and versatility of a variety of approaches for analyzing chromatin features in single cells. The improved data quality translates directly into better results for various types of downstream analysis. Accordingly, we anticipate that the methodological enhancement we have made will help to better resolve heterogeneity and variation of epigenetic gene regulation.

## Methods

### Tn5 loading for scTurboATAC

Lyophilized adapter oligonucleotides were resuspended in annealing buffer (50 mM NaCl, 40 mM Tris·HCl, pH 8) at 100 µM concentration. Adapters (**Table 1**) were preannealed (read 1 with the mosaic end (ME) sequence and read 2 with ME) separately on a thermocycler for 2 min at 85 °C followed by cooling down to 20 °C with 0.1 °C per minute. Glycerol was then added to a final concentration of 50% (v/v) and the adapters were stored at -20 °C. Purified unloaded Tn5 was obtained from the EMBL Protein Expression and Purification Core Facility, which was produced and stored as described previously (Hennig et al., 2018). For Tn5 assembly, transposase, annealed primer pairs and dilution buffer (DB, 50 mM Tris·HCl, pH 7.5, 100 mM NaCl, 0.1 mM EDTA, 1 mM DTT, 0.1 % NP-40, 50 % glycerol) were mixed with equal volumes and incubated for 30 min at room temperature. The loaded Tn5 was then diluted to 1:3 in DB yielding the Tn5-H100 preparation and stored at -20 °C.

### Comparing Tn5-H activity to Tn5-TXG

Activity was assesed by qPCR measuring the efficiency in generating fragmented lambda DNA. Different Tn5 preparations were used at the same volume with 10 ng of lambda DNA, 2.5 µl tagmentation buffer (20 mM Tris-HCl pH 7.5, 20 mM MgCl2, 50 % v/v DMF) or 10x Genomics tagmentation buffer as indicated. For the reaction, 1.25 µl phage DNA (10 ng), and 1.25 µl Tn5 enzyme were mixed and incubated for 10 min at 55 °C. The reaction was stopped by adding 1.25 µl 0.2 % SDS. The mix was diluted 1:100 in nuclease free water. A 10 µl qPCR mix was used that contained 5 µl 2x KAPA SYBR Fast, 0.2 µl 10 µM i7 adapter, 0.2 µl 10 µM i5 adapter, 1 µl diluted template DNA and 3.6 µl nuclease free water. The PCR program was 72 °C for 5 min, 95°C for 30 s, followed by 25 cycles of 95 °C for 10 s, 60 °C for 30 s and 72 °C for 60 s. Relative activities were calculated from the measured Ct values.

### scTurboATAC library preparation

For scTurboATAC, the transposase provided with the ATAC kit from 10x Genomics was replaced with Tn5-H100. Purified nuclei were processed using the standard 10x ATAC protocol according to the user guides for Chromium Next GEM Single Cell ATAC v1.1 or v2 kits (CG000496, 10x Genomics) as indicated. MEFs were cultured and treated as previously described (Muckenhuber et al., 2023). PBMCs were isolated as previously described and stored viably frozen in serum containing DMSO (Mallm et al., 2019). Prior to the experiments they were thawed in a water bath and immediately processed. For transposition, 10,000 nuclei were used and loaded on to the Chromium Next GEM Chip H (PN-1000161). Libraries were generated using standard reagents. Quality control and molarity calculations of the final libraries were performed with the Qubit 3.0 Fluorometer (Q33216, Invitrogen) and the 4200 tapestation system (Agilent Technologies). For multiome analysis the appropriately loaded Tn5 was used (**Table 1**), while all downstream reactions were performed with standard multiome reagents from 10x Genomics.

### scC&T & RNA from the same cell

TF1-IDH2 wildtype and mutant cell lines were acquired form ATCC (Manassas, Virginia, USA, wildtype, CRL-2003; IDH2 mutant, CRL-2003). They were cultured following the providers instructions. Cells were harvested, washed once in PBS and 1×10^6^ cells were fixed with 1 ml 0.2 % PFA for five min at room temperature. Fixation was stopped with 125 mM glycine for 5 min followed by one wash with PBS. Cells were then resuspended in 500 µl ice cold nuclei buffer and incubated for 10 min on ice. Cells were pelleted and resuspended in 500 µl antibody buffer (2 mM EDTA, 0.1 % BSA, 0.04 % digitonin, 20 mM HEPES pH 7.5, 150 mM NaCl and 0.5 mM spermidine).

A 10 µl volume of αH3K27me3 (cell signalling, 9733) and 12.5 µl RNase inhibitor (40 U/µl) were added and incubated over night at 4 °C. Cells were washed with wash buffer (0.04 % digitonin, 20 mM HEPES pH 7.5, 150 mM NaCl and 0.5 mM spermidine), resuspended in 500 µl wash buffer containing 10 µl 1 µg/µl anti-rabbit secondary antibody raised in guinea pig (Sigma-Aldrich, SAB3700889) and 1 U/µl RNase inhibitor. After 30 min at room temperature cells were washed three times with wash buffer with five min incubation time per wash and resuspended in Tn5 buffer (0.01 % digitonin, 20 mM HEPES pH 7.5, 150 mM NaCl, 0.5 mM spermidine and 1X protease inhibitor). Cells were counted and 500,000 cells were used from this point forward. 10 µl of loaded proteinA-Tn5 was added and incubated at room temperature for 30 min. After three washes with wash buffer, cells were resuspended in tagmentation buffer (10 mM Tris pH 7.5, 10 mM MgCl, 5 mM HEPES, 50 mM NaCl, 25 % DMF, 0.01 % digitonin, 1 U/µl RNase inhibitor), transferred to a PCR tube and incubated for one hour at 37 °C. A volume of 15 µl was then used for GEM formation with the multiome kit from 10x Genomics.

### Sequencing

Multiplexed library pools were generated at 10 nM concentrations of each library and sequenced on NovaSeq 6000 system (Illumina, v1.5). Sequencing was paired end with Read1N and Read2N sequenced with 50 cycles each, i7 (index) sequenced with 8 cycles and i5 using 10x Genomics barcodes with 16 cycles for ATAC and 24 cycles for multiome-ATAC and multiome-C&T. Sequencing depth was at least 25,000 read pairs per nucleus. RNA libraries were sequenced paired end with 28 bp and 90 bp for read 1 and read 2, respectively.

### Analysis of scATAC-seq data from MEFs

Data were demultiplexed and aligned with Cell Ranger ATAC count v2.0.0 (10x Genomics) using the provided mouse mm10 reference. Further processing of data was conducted with ArchR (v1.0.3) in R (v4.2.2) (Granja et al., 2021). Non-cell barcodes were removed by applying a minimal threshold to log10 number of fragments (3.8 for conventional scATAC, 3.9 for scATAC with Tn5-H30, and 4.3 for TurboATAC) and TSS enrichment score (5 for conventional ATAC, 12 for scATAC with Tn5-H30 and TurboATAC). This yielded 5153 cells for conventional scATAC, 5809 cells for scATAC with Tn5-H30, and 5612 cells for TurboATAC. Cell doublets were removed using Amulet (scDblFinder v1.12.0) (Thibodeau et al., 2021) and single cells were embedded in two-dimensional space using an accessibility matrix of 500 bp tiles, Iterative LSI (LSI components 2-10 for conventional ATAC, 2-12 for TurboATAC) and UMAP. Cells were clustered by SNN modularity optimization with a resolution of 0.1. Apoptotic cell clusters were identified by marker gene modules of single-cell gene activity scores (*Casp3, Parp1, Bax, Bak1, Bid, Bbc3, Pmaip1, Bcl2, Bcl2l1, Mcl1, Trp53, Cycs, Apaf1, Tnfrsf10b*, and *Fas*; (Galluzzi et al., 2018)) and removed for further analyses.

Sample-specific and merged peak sets were obtained from pseudo-bulk accessibility data using MACS2 (Zhang et al., 2008) in ArchR (extendSummits = 1000; reproducibility = 1). Transcription factor binding motifs were annotated in sample-specific peaks with Homer mm10 motifs from chromVARmotifs (v0.2.099) (Schep et al., 2017). Transcription factor motif footprints were computed using ArchR (no Tn5 bias normalization, 5 bp smooth window). MEF subtype clusters were identified as epithelial- and mesenchymal-like MEFs by marker gene modules of single-cell gene activity scores of epithelial- and mesenchymal-like MEF markers from our previous study (Muckenhuber et al., 2023).

Simultaneously accessible regions in the same cell were determined by co-accessibility analysis with RWire (https://github.com/RippeLab/RWire-IFN) based on our previously described RWire approach (Mallm et al., 2019;Muckenhuber et al., 2023). Cell numbers between the samples and subtypes were matched to 1,617 cells. The consensus peak set was used for all samples and subtypes. Pearson correlation coefficients were calculated for peaks within 1 Mb for each sample and subtype without aggregation of single cells. Only co-accessible links with positive correlation above background co-accessibility threshold r (0.088 for epithelial-like MEFs from conventional ATAC, 0.079 for mesenchymal-like MEFs from conventional ATAC, 0.066 for epithelial-like MEFs from TurboATAC, 0.066 for mesenchymal-like MEFs from TurboATAC) were considered. The background co-accessibility threshold was determined by the 99th co-accessibility percentile from randomly shuffled accessibility values over cells and peaks. Co-accessible links were further filtered by a lower threshold of 5 % accessible cells in the linked peaks. STAT1/2 bound sites and TSSs of interferon-stimulated genes from our previous study (Muckenhuber et al., 2023) at co-accessibly linked peaks were annotated.

### Analysis of scATAC-seq data from PBMCs

Data were demultiplexed and aligned with Cell Ranger ATAC count v2.1.0 (10x Genomics) using the provided human GRCh38 reference. Further processing of data was conducted with ArchR (v1.0.3) in R (v4.2.2) (Granja et al., 2021). Non-cell barcodes were removed by applying a minimal threshold to log10 number of fragments (4.1 for conventional ATAC, and 4.4 for TurboATAC) and TSS enrichment score (10 for conventional ATAC, 8 for TurboATAC). This yielded 7658 cells for conventional ATAC, and 8309 cells for TurboATAC. Cell doublets were removed using Amulet (scDblFinder v1.12.0) (Thibodeau et al., 2021) and single cells were embedded in two-dimensional space using an accessibility matrix of 500 bp tiles, Iterative LSI (LSI components 2-21) and UMAP. Cells were clustered by SNN modularity optimization with a resolution of 0.2.

Clusters were assigned to cell types by marker gene modules of single-cell gene activity scores of established markers (**Table S2**). Sample-specific and merged peak sets were obtained from pseudo-bulk accessibility data using MACS2 (Zhang et al., 2008) in ArchR (extendSummits = 1000; reproducibility = 1). Transcription factor binding motifs were annotated in sample-specific peaks with Homer hg38 motifs from chromVARmotifs (v0.2.0) (Schep et al., 2017). A subset of only B cells was embedded in two-dimensional space using an accessibility matrix of 500 bp tiles, Iterative LSI (LSI components 2-8) and UMAP. Cells were clustered by SNN modularity optimization with a resolution of 0.25. B cells from conventional ATAC and TurboATAC were integrated using Harmony (v0.1.1) (Korsunsky et al., 2019). ChromVAR deviations were calculated for 200 bp windows around top 1000 transcription factor binding motifs in sample-specific peak sets by decreasing scores (v1.20.0) (Schep et al., 2017). ChromVAR deviations were imputed using MAGIC (van Dijk et al., 2018) in ArchR.

### Analysis of scMultiome-ATAC data from PBMCs

Data were demultiplexed and aligned with Cell Ranger ARC count v2.0.2 (10x Genomics) using the provided human GRCh38 reference. Further processing of data was conducted with ArchR (v1.0.3) (Granja et al., 2021) and Seurat (v4.3.0) (Stuart et al., 2019) in R (v4.2.2). Non-cell barcodes in ATAC were removed by applying a minimal threshold of 3.8 and 1000 to log10 number of fragments and reads in TSS, respectively. This yielded 4563 cells for conventional ATAC, 7405 cells for ATAC Tn5-H50, and 6690 cells for TurboATAC. ATAC cell doublets were removed using Amulet (scDblFinder v1.12.0) (Thibodeau et al., 2021), cells with a blacklist ratio above 0.01 and/or nucleosome ratio above 3 were excluded, and single cells were embedded in two-dimensional space using an accessibility matrix of 500 bp tiles, Iterative LSI (LSI components 2-20) and UMAP. Cells were clustered by SNN modularity optimization with a resolution of 0.3.

Non-cell barcodes in RNA were removed by applying a minimal threshold of 100 and 500 to number of detected genes and number of UMI counts, respectively, as well as a maximal threshold of 40 % to percentage of mitochondrial UMI counts. This yielded 4587 cells for RNA of conventional ATAC, 7916 cells for RNA of ATAC Tn5-H50, and 7101 cells for RNA of TurboATAC. RNA cell doublets were removed using Scrublet with a cutoff of 0.15 (v0.2.3) (Wolock et al., 2019) in Python v3.10.4 (pandas v1.4.3, scipy v1.8.1) and single cells were embedded in two-dimensional space using SCTransform (v0.3.5) (Hafemeister and Satija, 2019), PCA (PCs 1-20) and UMAP. Cells were clustered by SNN modularity optimization with a resolution of 0.5. Co-embedding of single cells using ATAC and RNA information was conducted using Signac (v1.9.0) (Stuart et al., 2021). Combining filtered RNA and ATAC cell information yielded 3620 cells for RNA/ATAC of conventional ATAC, 5999 cells for RNA/ATAC of ATAC Tn5-H50, and 5077 cells for RNA/ATAC of TurboATAC. Single cells were embedded using LSI (LSI components 2-25) for ATAC, SCTransform and PCA (PCs 1-30) for RNA, weighted nearest neighbor graph and UMAP. Cells were clustered by SNN modularity optimization with a resolution of 0.8.

### Analysis of scMultiome-C&T (H3K27me3) data from TF1 cell lines

Data were demultiplexed and aligned with Cell Ranger ARC count v2.1.0 (10x Genomics) using the provided human GRCh38 reference. Further processing of data was conducted with ArchR (v1.0.3) (Granja et al., 2021) and Seurat (v4.3.0) (Stuart et al., 2019) in R (v4.2.2). Browser tracks of pseudo-bulk RNA and H3K27me3 signal were generated with plotgardener (v1.4.1) (Kramer et al., 2022). For H3K27me3 peak calling, MACS2 was used with the broad setting and keeping one read duplicate resulting in 57,533 peaks that were used in downstream analysis with ArchR. Non-cell barcodes in H3K27me3 were removed by applying a minimal threshold of 500 and 0.1 to number of fragments and fraction of reads in peaks, respectively. This yielded 2786 cells for H3K27me3 of TF1-IDH2wt, and 2653 cells for H3K27me3 of TF1-IDH2mut. Non-cell barcodes in RNA were removed by applying a minimal threshold of 100 and 500 to number of detected genes and number of UMI counts, respectively, as well as a maximal threshold of 25 % to percentage of mitochondrial UMI counts. This yielded 2928 cells for RNA of TF1-IDH2wt, and 2956 cells for RNA of TF1-IDH2mut. Co-embedding of single cells using H3K27me3 and RNA information was conducted using Signac (v1.9.0) (Stuart et al., 2021). Combining filtered RNA and H3K27me3 cell information yielded 1803 cells for RNA/H3K27me3 of TF1-IDH2wt, and 1800 cells for RNA/H3K27me3 of TF1-IDH2mut. Single cells were embedded using LSI (LSI components 3-20) for H3K27me3, SCTransform (v0.3.5) (Hafemeister and Satija, 2019) and PCA (PCs 1-20) for RNA, weighted nearest neighbor graph and UMAP.

## Data availability

Single cell sequencing data are available via GEO accession number GSE235506.

## Acknowledgments

We thank Markus Muckenhuber for discussions and Sabrina Schumacher for technical assistance. This work was supported by German Research Foundation (DFG) grant FOR2674 (Z01) to JPM and KR and grants TRR179 (Z03) and SFB1074 (Z01) to KR and by the German Federal Ministry of Education and Research (BMBF) via project SATURN3 (01KD2206B) within the National Decade against Cancer program to KR. We are grateful to the EMBL Protein Expression and Purification Core Facility for providing the Tn5 protein preparation. We thank the DKFZ Genomics and Proteomics and Omics IT and Data Management Core Facilities for sequencing and data management.

## Notes

### Competing Interest Statement

The authors have declared no competing interest.

