## Supplementary information for "Enhancing sensitivity and versatility of Tn5-based single cell omics"

### Inventory of Supplementary Information

#### Supplementary Figures

Figure S1. Tn5 activity measurements by qPCR

Figure S2. Quality assessment of scTurboATAC data

Figure S3. Investigation of transcription regulation with scTurboATAC

Figure S4. Resolving cell types in PBMCs with scTurboATAC

Figure S5. Quality assessment of single cell multiome data

#### Supplementary Tables

Table S1. Sequencing metrics of scATAC and scTurboATAC analysis

Table S2. Marker genes of hematopoietic cell types

Table S3. Sequencing metrics of scMultiome-ATAC in PBMCs

Table S4. Sequencing metrics of scMultiome-C&T in TF1 cell lines

#### Supplementary References

### Supplementary Figures

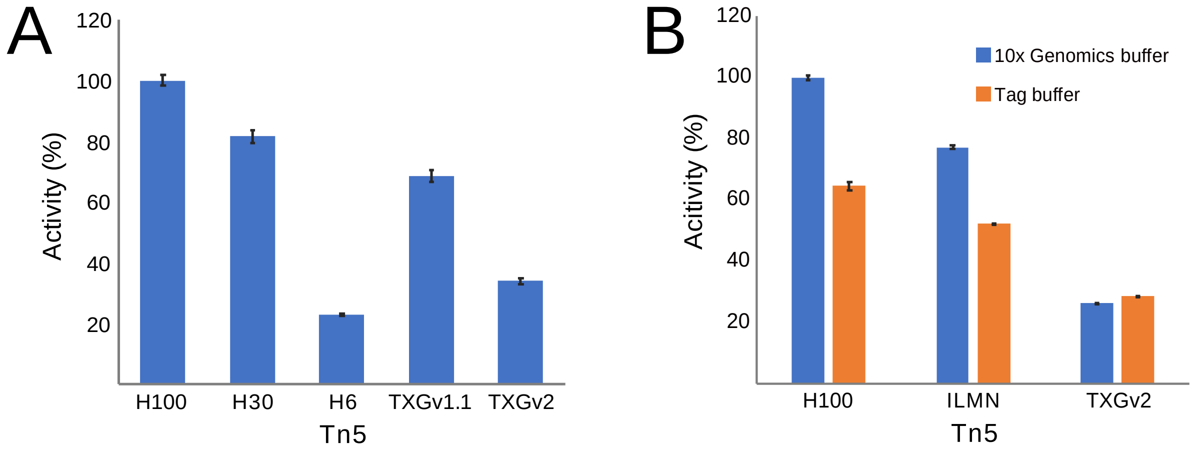


**Figure S1. Tn5 activity measurements by qPCR.** (**A**) The in-house Tn5-H preparations at three different concentrations (Tn5-H100, Tn5-H30 and Tn5-H6) were tested against Tn5-TXGv1.1 and Tn5-TXGv2 that are provided in the kits from 10x Genomics. Activity was calculated from the yield of fragmented lambda phage DNA measured by qPCR. Tn5-H100 displayed the highest activity while Tn5-TXGv1.1 was more efficient than the later introduced Tn5-TXGv2 enzyme. Error bars display standard deviation from triplicates. (**B**) Comparison of Tn5 activity in the buffer from 10x Genomics against a standard Tn5 reaction buffer (Tag buffer) for the different Tn5 enzyme preparations. The buffer provided in the 10x Genomics kits results in a higher activity for both the Tn5-H100 as well as the Tn5-ILMN (TDE1 enzyme, #20034197, Illumina) preparations. No difference between the two buffers was observed for Tn5-TXGv2. Error bars display standard deviation from triplicates.

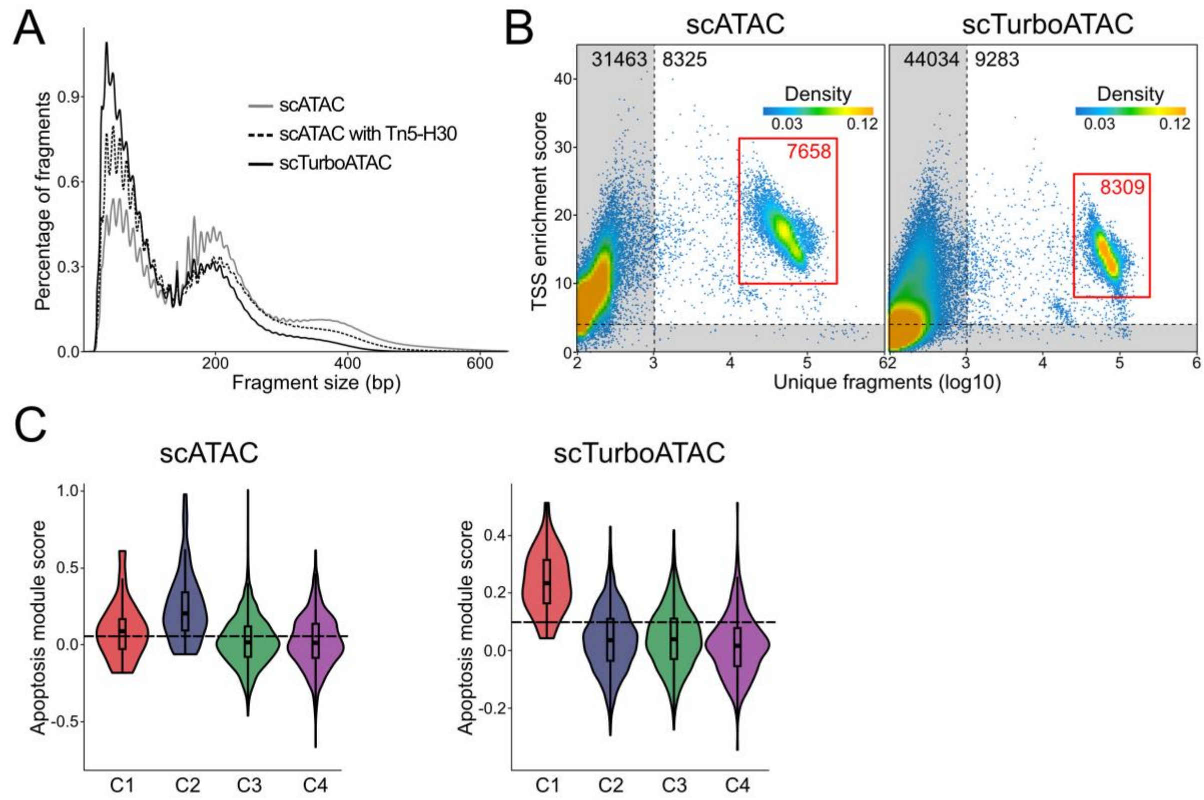


**Figure S1. Quality assessment of scTurboATAC data.** (**A**) Fragment size distribution in scATAC with Tn5-TXGv1.1 and Tn5-H30 versus scTurboATAC protocols for MEFs stimulated for 6h with IFNβ. (**B**) TSS enrichment score against log10 number of unique fragments of cell barcodes in scATAC with Tn5-TXGv2 (left) and scTurboATAC (right) for PBMCs. Colors of points reflect density of cell barcodes. Grey area marks low-quality cell barcodes. Red rectangle marks selected high-quality cells. (**C**) Apoptosis module score per cell in scATAC with Tn5-TXGv1.1 (left) and scTurboATAC (right) of 6h IFNβ-stimulated MEFs.

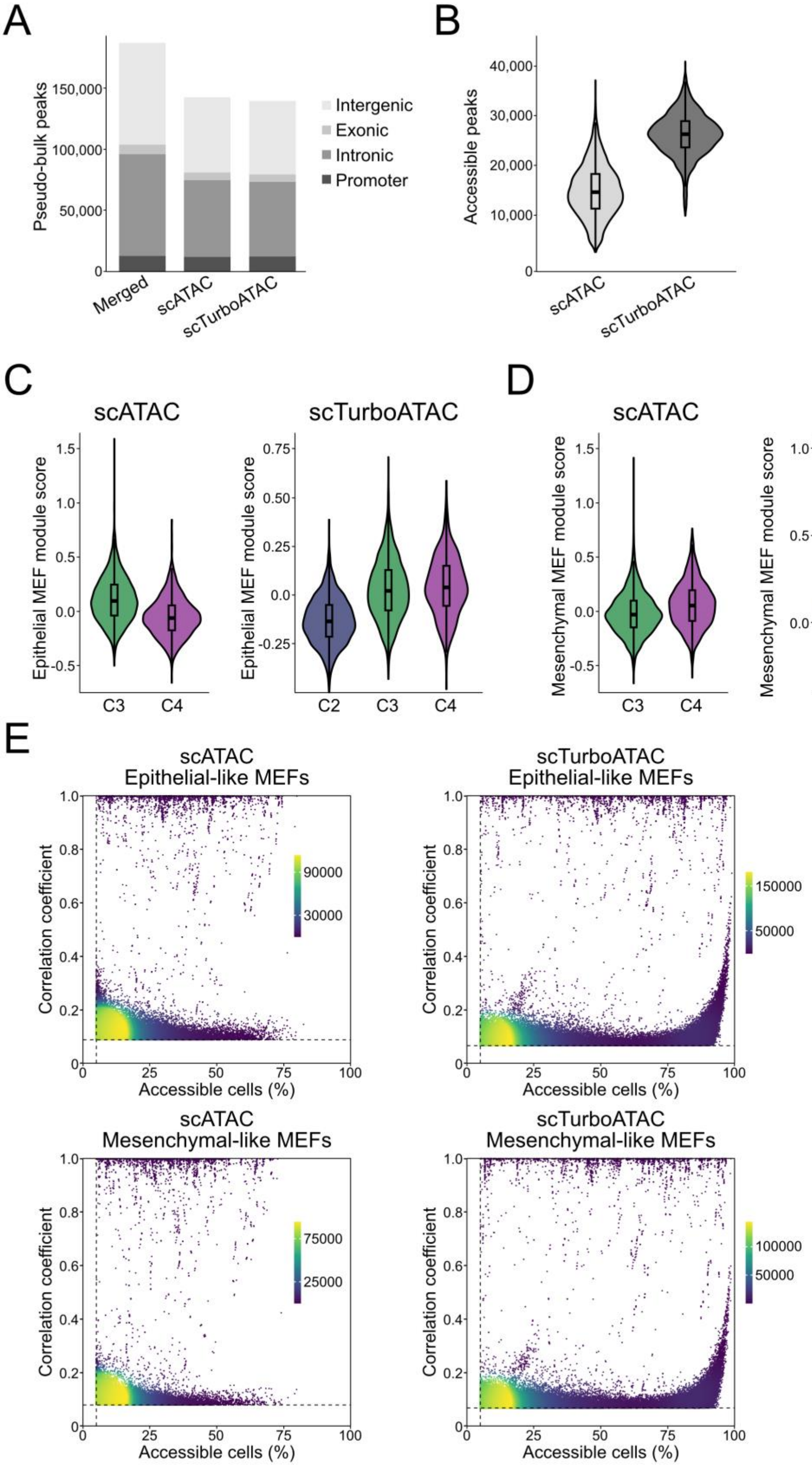


**Figure S2. Investigation of transcription regulation with scTurboATAC.** MEFs stimulated for 6h with IFNβ were studied. The scATAC data refer to the Tn5-TXGv1.1 enzyme. (**A**)

Number of peaks from pseudo-bulk chromatin accessibility of scATAC, scTurboATAC and the merged peak set, which additionally included data from scATAC with Tn5-H30. (**B**) Number of accessible peaks detected per cell in scATAC and scTurboATAC using the merged peak set. (**C**) Epithelial-like MEF module score per cell in scATAC and scTurboATAC. (**D**) Mesenchymal-like MEF module score per cell in scATAC and scTurboATAC. (**E**) Correlation coefficient and percent accessible cells of co-accessible links from scATAC (left) and scTurboATAC (right). Colors of points reflect density of co-accessible links. Dashed lines represent background co-accessibility cutoff on correlation coefficient and minimal 5 % cutoff on accessible cells.

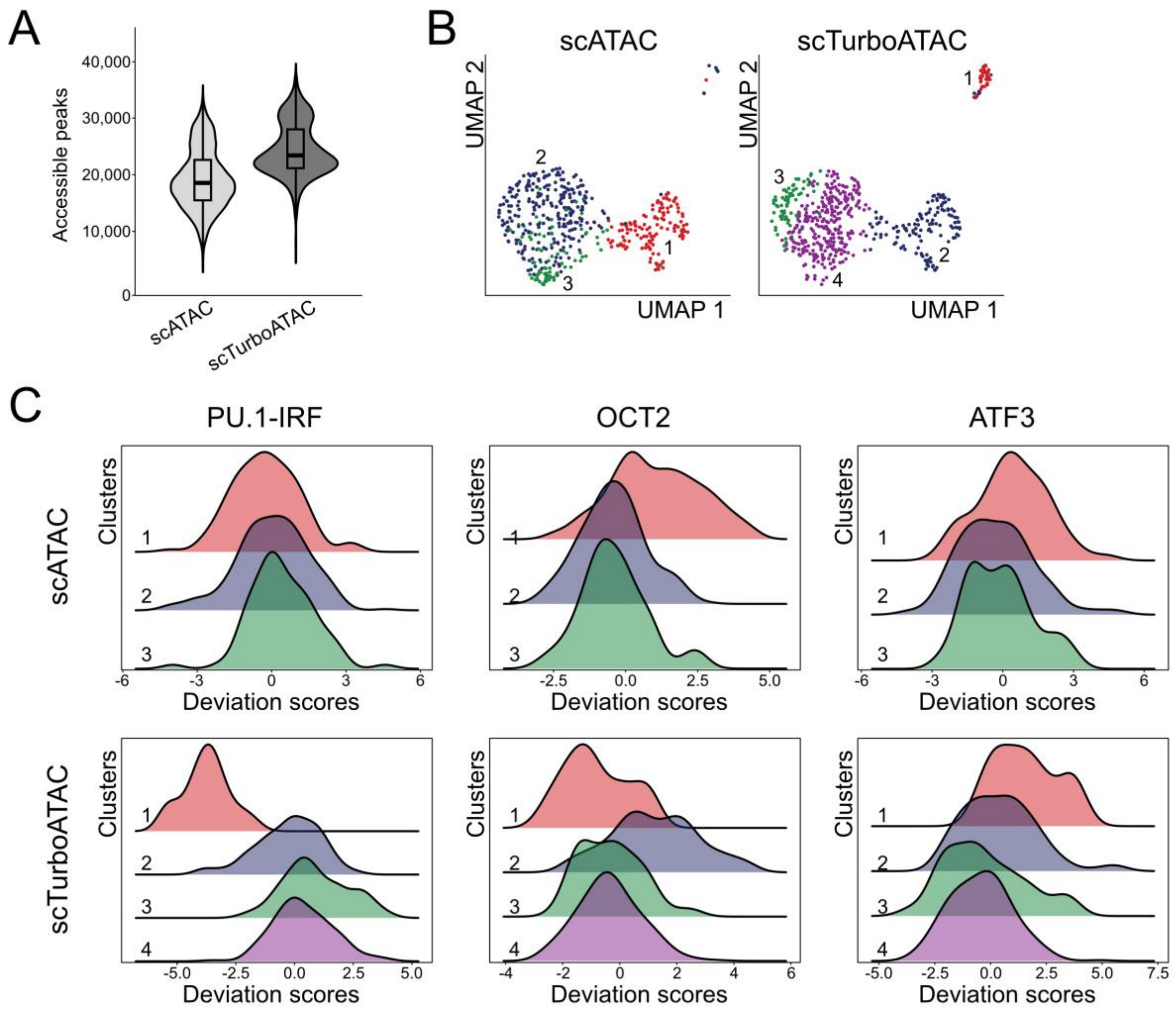


**Figure S3. Resolving cell types in PBMCs with scTurboATAC.** The scATAC data refer to the Tn5-TXGv2 enzyme. (**A**) Number of accessible peaks per cell in scATAC and scTurboATAC using the merged peak set. (**B**) Integrated UMAP embedding of B cells filtered from PBMC data set for scATAC and scTurboATAC. Integrated UMAP embeddings split by samples scATAC (left) and scTurboATAC (right) are shown. Cells are colored according to sample-specific k-nearest neighbor clusters. (**C**) Transcription factor motif deviation scores (non-imputed) in k-nearest neighbor clusters of scATAC (top) and scTurboATAC (bottom).

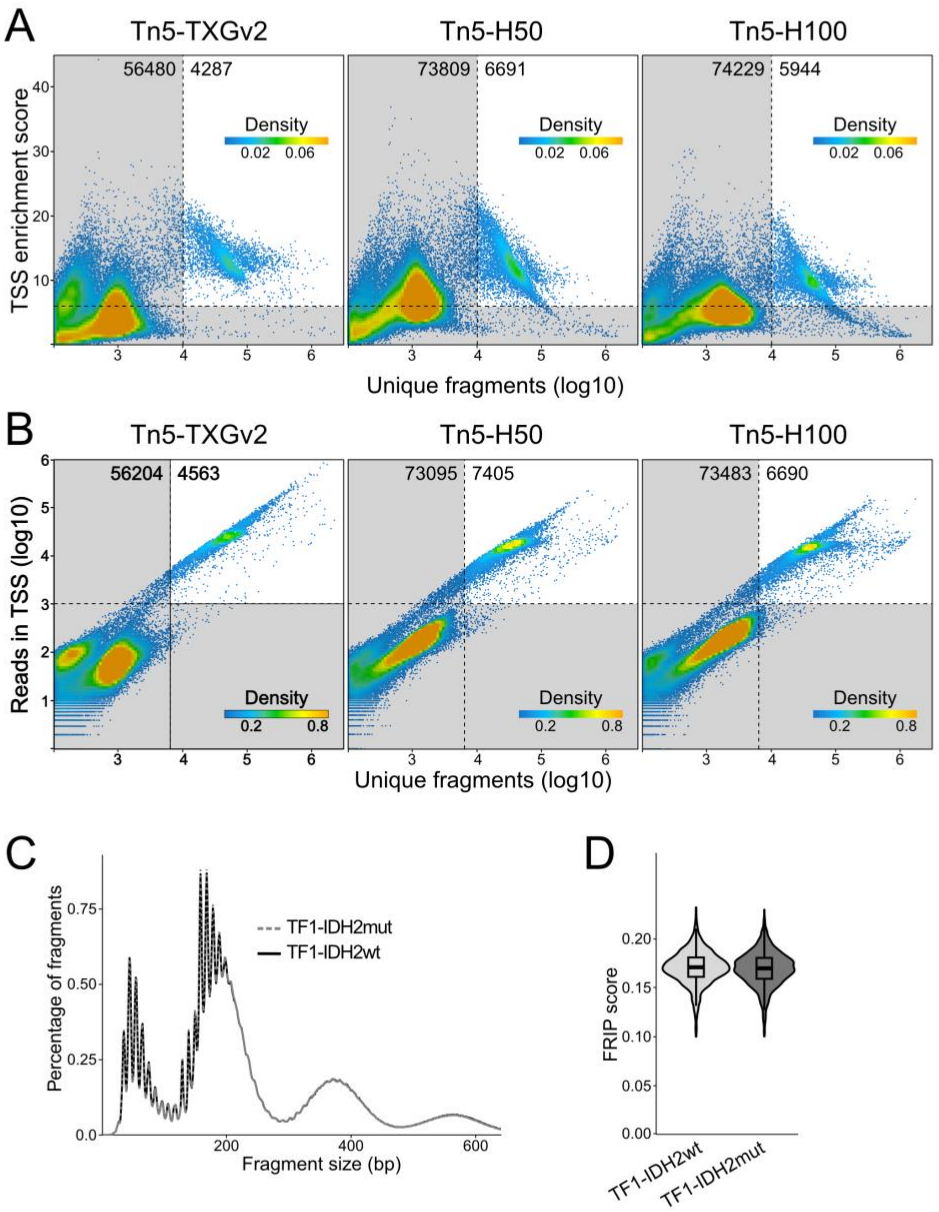


**Figure S5. Quality assessment of single cell multiome data.** (**A**) TSS enrichment score over log10 number of unique fragments of cell barcodes in scMultiome-ATAC of PBMCs with the indicated Tn5 preparations. Colors of points reflect density of cell barcodes. Grey area marks low-quality cell barcodes. (**B**) Same as panel A for log10 number of reads in TSS over log10 number of unique fragments. (**C**) Fragment size distribution for scMultiome-C&T against H3K27me3 in TF1-IDH2wt and TF1-IDH2mut. (**D**) Fragments of reads in peaks (FRIP) scores for H3K27me3 per cell in TF1-IDH2wt and TF1-IDH2mut from the scMultiome-C&T analysis.

### Supplementary Tables

#### Table S1. Sequencing metrics of scATAC and scTurboATAC analysis

| | **MEF 6h IFNβ** | | | **PBMC** | |
|---|---|---|---|---|---|
| | scATAC with Tn5-TXGv1.1 | scATAC with Tn5-H30 | scTurboATAC | scATAC with Tn5-TXGv2 | scTurboATAC |
| Sequenced read pairs | 827,356,762 | 1,221,969,310 | 1,304,430,940 | 1,466,934,091 | 1,468,516,560 |
| Duplicates | 70.1 % | 75.8 % | 45.9 % | 66.6 % | 52.2 % |
| Mapped read pairs | 92.2 % | 88.2 % | 87.7 % | 94.0 % | 90.5 % |
| Fragments nucleosome-free regions | 40.6 % | 54.5 % | 64.8 % | 50.4 % | 47.5 % |

**Table S2. Marker genes of hematopoietic cell types**

| Cell type | Marker gene | Reference |
|---|---|---|
| Naïve CD8+ T cells | CD8A, GZMB, CD3D, CD34, LEF1 | (Terstappen et al., 1992; Uhlen et al., 2019) |
| Mature CD8+ T cells | CD8A, GZMB, CD3D, CD2 | (Uhlen et al., 2019) |
| Naïve CD4+ T cells | CD4, IL7R, CCR7, CD3D, CD34, LEF1 | (Terstappen et al., 1992; Uhlen et al., 2019) |
| Mature CD4+ T cells | CD4,IL7R, S100A4, CD3D, CD2, IL2, TNF, IL21 | (Uhlen et al., 2019) |
| NK cells | GNLY, NKG7, GZMB, NCAM1, FCGR3A | (Uhlen et al., 2019; Karlsson et al., 2021) |
| B cells | MS4A1, CD79A, CD74, CD19 | (Uhlen et al., 2019; Karlsson et al., 2021) |
| Progenitor cells | CD34, KIT, PROM1, PTPRC (negative marker), CD38 (negative marker) | (Terstappen et al., 1992; Monaco et al., 2019) |
| Classical monocytes | CD14, LYZ, CST3 | (Uhlen et al., 2019) |
| Non-classical monocytes | FCGR3A, MS4A7 | (Uhlen et al., 2019) |
| Dentritic cells | FCER1A, CST3, CD74 | (Uhlen et al., 2019) |
| Basophils | ENPP3, CD69, CCR3, PTGDR2 | (Monaco et al., 2019; Uhlen et al., 2019) |
| Neutrophils | CEACAM8, ITGAM, FCGR3B, PTPRC | (Uhlen et al., 2019)(Monaco et al., 2019) |
| Eosinophils | EPX, RNASE2, CLC, SIGLEC8 | (Uhlen et al., 2019) |
| Erythrocytes | HBA2, HBB, HBA1, SLC4A1, EPB41 | (Karlsson et al., 2021) |
| Platelets | PPBP, ITGA2B, PF4 | (Shattil et al., 1985; Poncz et al., 1987; Majumdar et al., 1991) |

**Table S3. Sequencing metrics of scMultiome-ATAC analysis in PBMCs**

| | **Tn5-TXGv2** | **Tn5-H50** | **Tn5-H100** |
|---|---|---|---|
| Sequenced ATAC read pairs | 1,304,897,617 | 1,431,221,793 | 1,444,079,941 |
| Sequenced RNA read pairs | 714,778,267 | 644,951,970 | 716,455,503 |
| Duplicates ATAC | 68.5 % | 67.2 % | 57.3 % |
| Duplicates RNA | 92.3 % | 91.8 % | 92.2 % |
| Mapped ATAC read pairs | 92.5 % | 91.0 % | 91.2 % |
| Mapped RNA read pairs | 61.4 % | 53.1 % | 51.8 % |

**Table S4. Sequencing metrics of scMultiome-C&T in TF1 cell lines**

| | **TF1-IDH2wt** | **TF1-IDH2mut** |
|---|---|---|
| Sequenced H3K27me3 read pairs | 486,259,010 | 468,044,936 |
| Sequenced RNA read pairs | 42,673,800 | 36,916,878 |
| Duplicates H3K27me3 | 96.6 % | 96.6 % |
| Duplicates RNA | 27.0 % | 29.5 % |
| Mapped H3K27me3 read pairs | 84.1 % | 83.5 % |
| Mapped RNA read pairs | 68.5 % | 68.5 % |
